## Supplementary data for "Muscle fiber Myc is dispensable for muscle growth and forced expression severely perturbs homeostasis"

**SUPPLEMENTARY FIGURES**

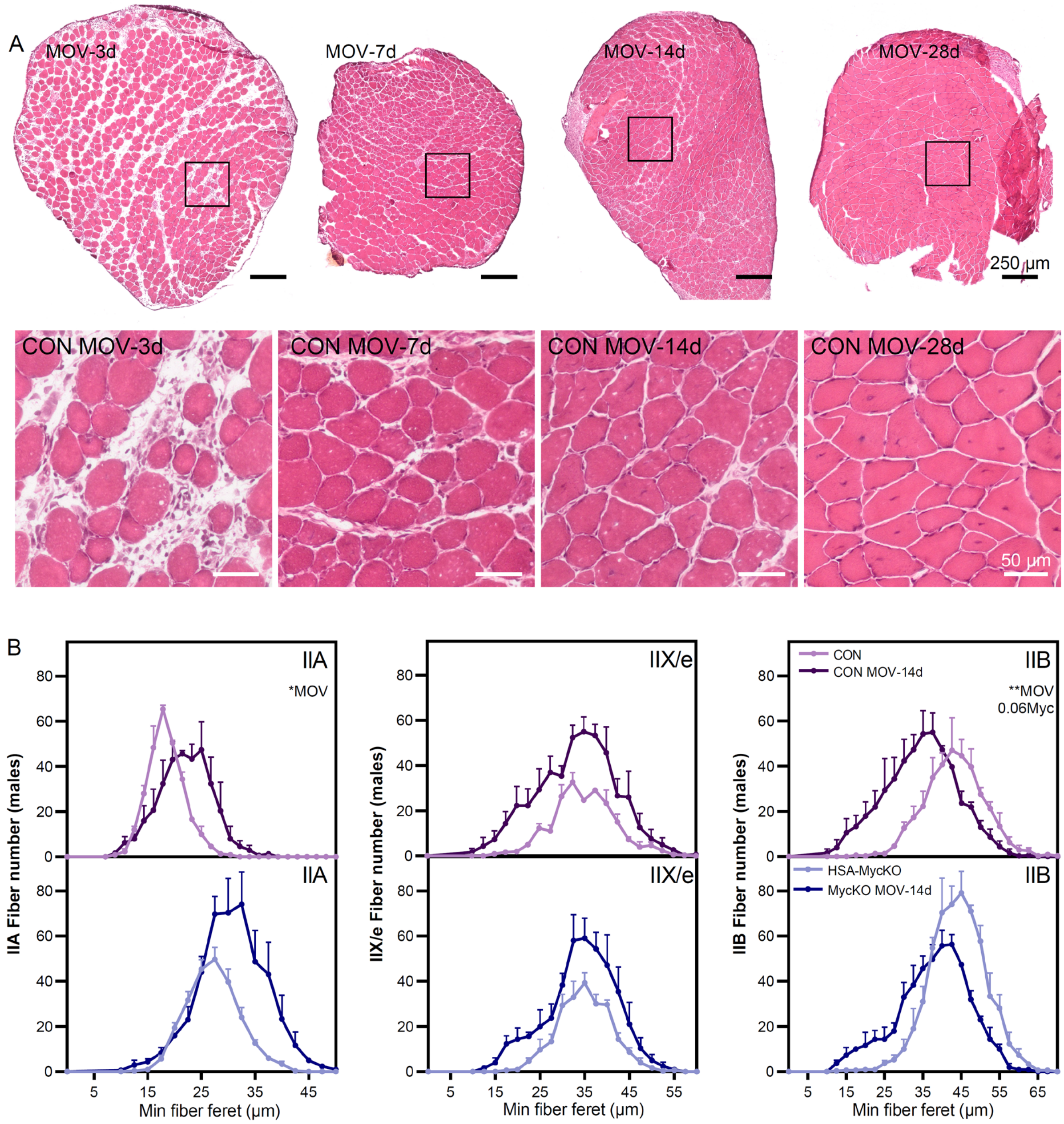


**Supplementary Figure 1. (A)** Representative haematoxolyin and eosin (H&E) images of *plantaris* (PLA) muscle cross-sections from CON mice after 3, 7, 14 and 28 days of mechanical overload (MOV). **(B)** Type-specific (IIA, IIX, IIB) fiber size distribution normalized to contralateral total fiber number after 14d of MOV in CON and HSA-MycKO mice. Data are presented as mean ± SEM. Two-way ANOVAs with Sidak's post hoc test were used to compare median fiber size between conditions. *, **, and *** denote a significant difference between groups of $P < 0.05$, $P < 0.01$, and $P < 0.001$, respectively.

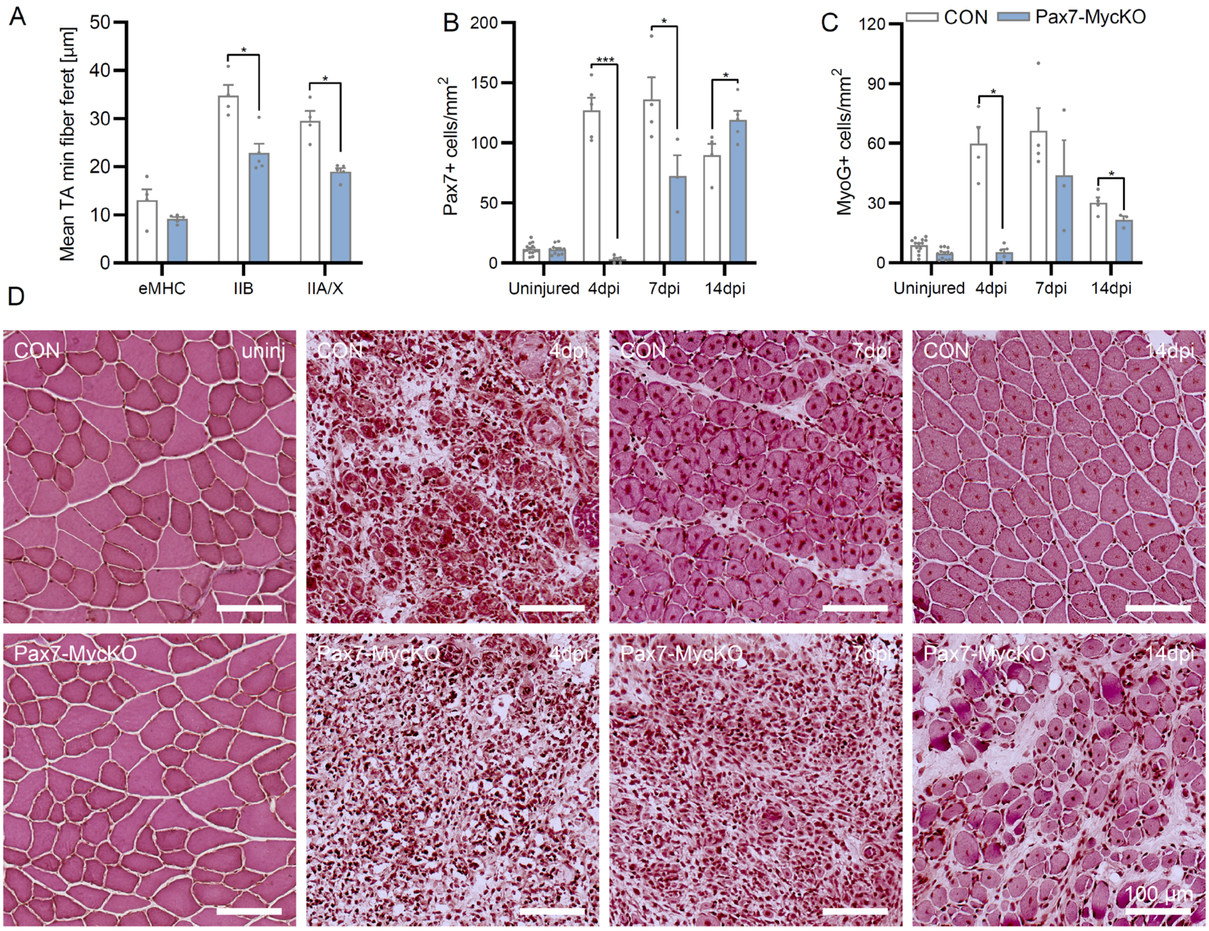


**Supplementary Figure 2. (A)** Mean minimum fiber feret size for eMHC, IIB and unstained fibers (IIA/X) 14 days after cardiotoxin (CTX) injury in CON and Pax7-MycKO TA cross-sections. Quantification of **(B)** Pax7+ and **(C)** MyoG+ cell number in TA muscle sections, via immunostaining, 4, 7 and 14 days after CTX injury in CON and Pax7-MycKO mice normalized to cross-sectional area. Uninjured samples represent pooled, un-injected contralateral legs from all time points. **(D)** Representative haematoxolyin and eosin (H&E) images of TA muscle cross-sections from CON and Pax7-MycKO mice in uninjured muscle and 4, 7 and 14 days after cardiotoxin injection. Data are presented as mean ± SEM. Two-way ANOVAs with Sidak's post hoc test were used to compare between conditions. *, **, and *** denote a significant difference between groups of $P < 0.05$, $P < 0.01$, and $P < 0.001$, respectively.

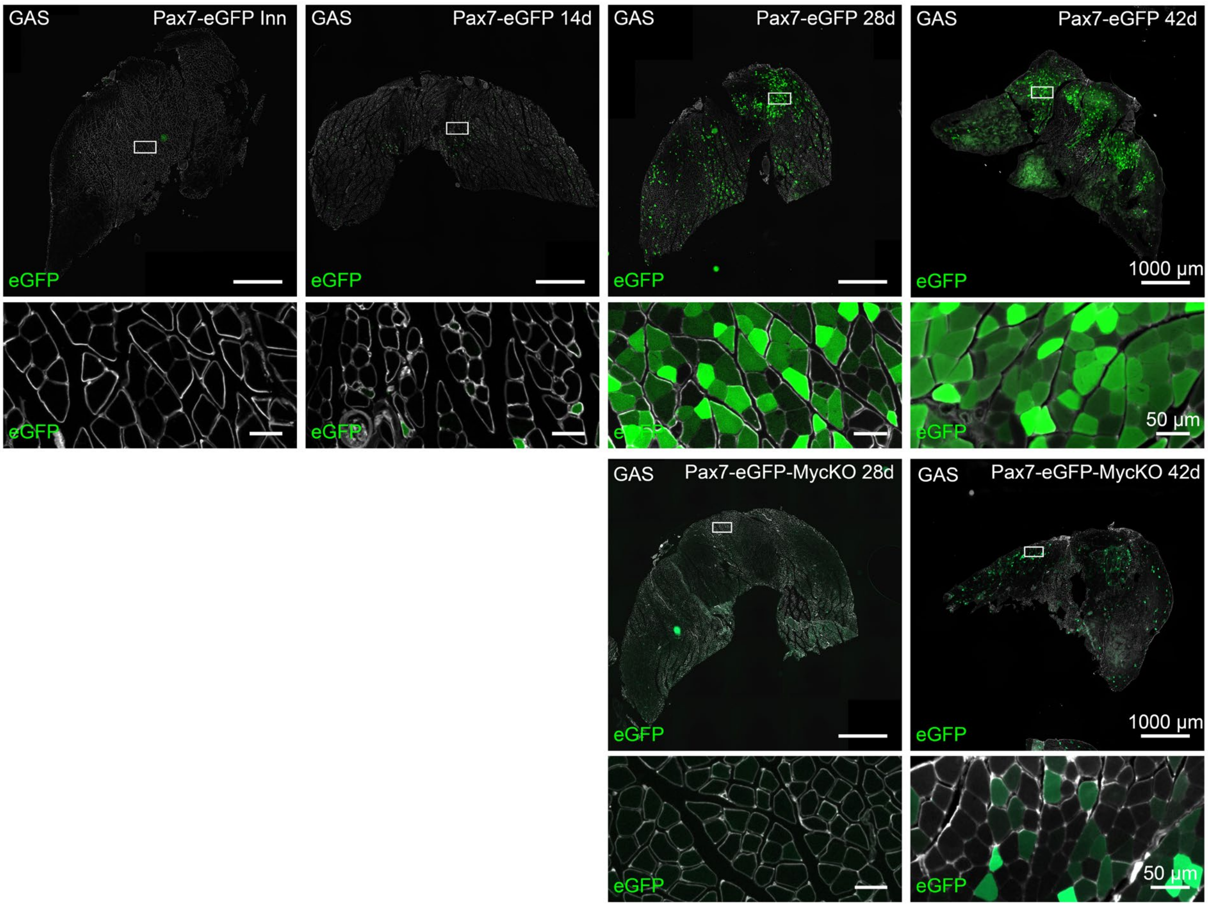


**Supplementary Figure 3.** Representative images of eGFP+ fibers, indicating fusion of Pax7-eGFP MuSCs in GAS muscle fibers ~6 weeks after tamoxifen treatment in contralateral control muscle (0d) and 14, 28 and 42 days after sciatic nerve crush in Pax7-eGFP mice and 28 and 42 days after nerve crush in Pax7-MycKO mice. Cross-sections are counterstained with laminin (white).

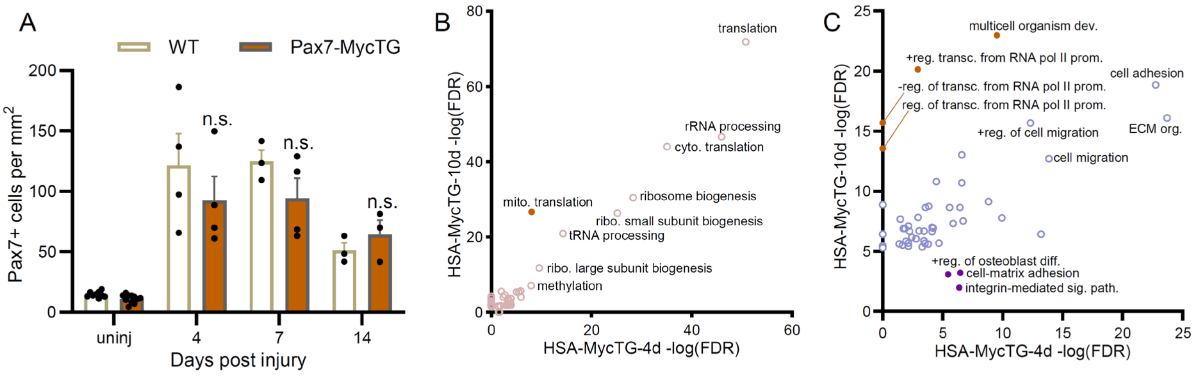


**Supplementary Figure 4. (A)** Pax7+ cell number normalized to cross-sectional area in TA muscle sections, in uninjured muscle and 4, 7 and 14 days after cardiotoxin (CTX) injection in CON and Pax7-MycTG mice. **(B)** Upregulated and **(C)** downregulated (right) gene ontology (GO) terms (Biological process) enriched in either 4d (x-axis) or 10d (y-axis) HSA-MycTG mice compared to CON mice. Top GO terms are labelled, while terms more prominently represented in 4d and 10d HSA-MycTG are shown in purple and orange, respectively. Data are presented as mean ± SEM. Two-way ANOVAs with Sidak's post hoc test were used to compare between conditions.
